# A cross-species approach using an *in vivo* evaluation platform in mice demonstrates that sequence variation in the human *RABEP2* gene modulates ischemic stroke outcomes

**DOI:** 10.1101/2022.06.29.498109

**Authors:** Han Kyu Lee, Do Hoon Kwon, Douglas A. Marchuk

## Abstract

Ischemic stroke, caused by vessel blockage, results in cerebral infarction; the death of brain tissue. Previously, quantitative trait locus mapping (QTL) of cerebral infarct volume and collateral vessel number identified a single, strong genetic locus regulating both phenotypes. Additional studies identified the causative gene, encoding RAB GTPase Binding Effector Protein 2 (*Rabep2*). However, there is yet no evidence that variation in the human ortholog of this gene plays any role in ischemic stroke outcomes. We established an *in vivo* evaluation platform in mice using adeno-associated virus (AAV) gene replacement and verified that both mouse and human RABEP2 rescue the mouse *Rabep2* KO ischemic stroke volume and collateral vessel phenotypes. Importantly, this cross-species complementation enabled us to experimentally investigate the functional effects of coding sequence variation in the human *RABEP2* gene. We chose four coding variants from the human population that are predicted by multiple *in silico* algorithms to be damaging to RABEP2 function. *In vitro* and *in vivo* analyses verify that all four led to decreased collateral vessel connections and increased infarct volume. Thus, there are naturally occurring loss-of-function alleles. This cross-species approach will expand the number of targets for therapeutics development for ischemic stroke.

## Introduction

Stroke is the second-leading cause of death in the world, with fifteen million new cases reported annually. In the United States, stroke is the fourth-leading cause of death and the primary cause of severe, long-term disability, with almost 800,000 new or recurrent cases occurring each year (1-5). Over 87% of strokes are ischemic, caused by a disruption of the blood supply to the brain leading to neuronal tissue damage (including infarction) in the ischemic region (6-8). Once a stroke has occurred, the only FDA-approved drug for stroke therapy is intravenous tissue plasminogen activator (IV tPA), currently given to only 2-3% of stroke patients because of its limited time window for administration and the major risk for adverse effects (9-13). Therefore, there is an urgent need to identify and develop new drug targets to effectively treat stroke patients. Decades of studies have identified individual risk factors for stroke (e.g., hypertension, smoking, diabetes, obesity, etc.) (14-17). Genetic risk factors for stroke susceptibility have been identified using both family-based linkage (18,19) and population-based genome-wide association studies (GWAS) (20-22). However, there have been no new targets identified by GWAS for stroke treatment.

In principle, new therapeutic targets could be identified by genetic studies of patients stroke outcomes and infarct volume (size). However, genetic mapping in the human population for stroke infarct volume is intrinsically problematic due to uncontrollable variation in the extent and location of the occluded vessel, and especially, variation in the critical time between the first recognized symptoms of stroke and medical intervention.

By contrast, the surgical occlusion mouse model of cerebral ischemia enables complete control over the variables that cannot be controlled in human patients. Thus, to discover novel genes modulating ischemic stroke infarction, we and others have taken a QTL (i.e., forward genetic mapping) analysis approach using the surgical occlusion mouse model of cerebral ischemia. Through permanent occlusion of the distal middle cerebral artery (pMCAO) in 32 commonly used inbred mouse strains, we have shown that these strains demonstrate robust differences in infarct volume. The differences in infarct volume are highly reproducible within each strain and differ more than 50-fold across all strains (23,24). Importantly, the differences are caused by natural allelic variation in the mouse genome. We have exploited these differences to map the natural genetic determinants of infarct volume for several genetic loci that modulate ischemic stroke, and have identified several causative genes (23-29). The strongest and most significant locus modulating infarct volume is found in multiple pairwise crosses of inbred mouse strains. This locus, cerebral infarct volume QTL 1 (*Civq1*), overlaps another locus on chromosome 7 which modulates cerebral collateral vessel number (collateral artery number QTL 1; *Canq1*) (30,31). The gene encoding *Rabep2*, an endosomal recycling protein for vascular endothelial growth factor receptor-2 (VEGFR2), was subsequently identified as the *Civq1/Canq1* gene (32,33).

However, the role of the human *RABEP2* gene in ischemic stroke remains uncertain. To date, no GWAS studies of ischemic stroke phenotypes have identified SNPs within or near this gene. This could be due to the aforementioned difficulty of controlling all of the non-genetic variables in stroke outcomes. Coding variations do exist in the human *RABEP2* gene and some of these variants, while rare overall in the human population, are predicted to be damaging to its function by *in silico* gene variant prediction algorithms. However, their effects have not been evaluated for ischemic stroke outcomes.

In the present study, we developed an *in vivo* evaluation platform to investigate the role of *Rabep2* on ischemic stroke volume. This platform also enabled us to examine the human *RABEP2* gene in *Rabep2* KO mice. We then evaluated coding variants of the human *RABEP2* gene, focusing on four of the most common variants that are also computationally predicted to abrogate its function.

## Material and methods

### Animals

*Rabep2* KO mice were generated by the Duke Transgenic Core using CRISPR/Cas9 technology to remove sequence from Exon3 of the gene, which was chosen to alter the reading frame downstream of the deletion. Briefly, CRISPR reagents including single-guide RNA and Cas9 protein were injected into C57BL6/J mouse by electroporation. Once founders were born, sequencing analysis for the targeted locus was performed to screen *Rabep2* KO mouse candidates that were maintained on a C57BL6/J background. Mice (both males and females) were age-matched (P21 and 10-week for collateral vessel perfusion and 12 ± 1 week for pMCAO) for all experiments.

### Study approval

All animal study procedures were conducted under a protocol approved by the Duke University IACUC in accordance with NIH guidelines.

### Viruses

All the AAVs used for this study were generated by the Duke Viral Vector Core. A control virus with CMV promoter (pAAV9-CMV-EGFP-WPRE) and six target viruses with CMV promoter containing either mouse *Rabep2* (pAAV9-CMV-m*Rabep2*-P2A-EGFP-WPRE), human *RABEP2* (pAAV9-CMV-h*RABEP2*-P2A-EGFP-WPRE), or four human RABEP2 coding variants (pAAV9-CMV-h*RABEP2*(Variants)-P2A-EGFP-WPRE; R508S, S204L, R490W, and R543H were used for experiments. Briefly, 1 μL of AAV construct (minimum titer, 2 × 10^13^ vg/mL) was directly injected into the right hemisphere of the brain of P1 *Rabep2* KO animals.

### Collateral vessel density measurement

As we have shown that collateral vessel traits are established by 3-week of age and remain constant for many months (25), the collateral vessel phenotype was measured at P21 and 10-week as previously described (26-29). Mice were anesthetized with ketamine (100 mg/kg) and xylazine (5 mg/kg), and the ascending thoracic aorta was cannulated. The animals were perfused with freshly made buffer (1 mg/mL adenosine, 40 g/ml papaverins, and 25 mg/mL heparin in PBS) to remove the blood. The pial circulation was then exposed after removal of the dorsal calvarium and adherent dura mater. The cardiac left ventricle was cannulated and a polyurethane solution with a viscosity sufficient to minimize capillary transit (1:1 resin to 2-butanone, PU4ii, VasQtec) was slowly infused; cerebral circulation was visualized under a stereomicroscope during infusion. The brain surface was topically rinsed with 10% PBS-buffered formalin and the dye solidified within 20 min. After post-fixation with 10% PBS-buffered formalin, pial circulation was imaged. All collateral interconnecting the anterior- and middle cerebral artery trees of both hemispheres were counted.

### pMCAO

Focal cerebral ischemia was induced by direct permanent occlusion of the distal MCA as previously described (26-29). Briefly, adult mice were anesthetized with ketamine (100 mg/kg) and xylazine (5 mg/kg), and then 0.5% bupivacaine (5 mg/mL) was also administrated by injection at the incision site. The right MCA was exposed by a 0.5 cm vertical skin incision midway between the right eye and ear under a dissecting microscope. After the temporalis muscle was split, a 2 mm burr hole was made with a high-speed micro drill at the junction of the zygomatic arch and the squamous born through the outer surface of the semi-translucent skull. The MCA was clearly visible at the level of the inferior cerebral vein. The inner layer of the skull was removed with fine forceps, and the dura was opened with 32-gauge needle. While visualizing under an operating microscope, the right MCA was electrocauterized. The cauterized MCA segment was then transected with microscissors to verify permanent occlusion. The surgical site was closed with 6-0 sterile nylon sutures, and 0.5 % bupivacaine was applied. The temperature of each mouse was maintained at 37°C with a heating pad during the surgery and then mouse was placed in a recovery chamber (set temperature 37°C) until the animal was fully recovered from the anesthetic. Mice were then returned to their cages and allowed free access to food and water in an air-ventilated room with the ambient temperature set to 25°C.

### Infarct volume measurement

Cerebral infarct volumes were measured 24 hours after distal permanent MCA occlusion, the time point when the size of the cortical infarct is largest and is stable. Twenty-four hours after pMCAO surgery, the animals were euthanized, and the brains were carefully removed. The brains were placed in a brain matrix, chilled at -80°C for 4 min to slightly harden the tissue, and then sliced into 1 mm coronal sections. Each brain slice was placed in 1 well of a 24-well plate and incubated for 20 min in a solution of 2% 2,3,5-triphenyltetrazolium chloride (TTC) in PBS at 37°C in the dark. The sections were then washed once with PBS and fixed with 10% PBS-buffered formalin at 4°C. Then 24 hours after fixation, the caudal face of each section was scanned using a flatbed color scanner. The scanned images were used to determine infarct volume. Image-Pro software (Media Cybernetics, Inc., MD) was used to calculate the infarcted area of the hemisphere to minimize error introduced by edema. The total infarct volume was calculated by summing the individual slices from each animal.

### Immunohistochemistry (IHC)

IHC was performed by Servicebio (Servicebio, Inc., MA). *Rabep2* KO mice either 3 weeks or 10 weeks after CMV-m*Rabep2* injection were fixed with 4% PFA and then mouse brains were shipped for sample preparations. Briefly, brain samples were processed, embedded in paraffin, and sagittally sectioned at 4μm. Paraffin sections were then deparaffinized with ethanol and rehydrated gradually with distilled water at room temperature. The sections were submerged in 0.01M sodium citrate buffer and boiled for 10 min for retrieval of antigen. The sections were washed with PBS three times and treated 3% H_2_O_2_ for 15 min and blocked with blocking solution (5% BSA) for 1 hour at room temperature before application of primary antibody. The sections were incubated with rabbit anti-GFP (1:1000, Cell Signaling Technology, MA) overnight at 4°C. Subsequently, the sections were immunohistochemically stained with Alexa Fluor 488 (1:1000, Molecular Probes, OR) for 1 hour at room temperature. Whole slide scanning was performed on a Panorama Midi II scanner (3DHISTECH, Ltd., Hungary) and images were captured using CaseViewer software (CaseViewer 2.3, 3DHISTECH, Ltd., Hungary).

### *In silico* prediction algorithms

Coding polymorphisms were examined for potential functional consequences using two independent *in silico* prediction algorithms, Polymorphism Phenotyping v2 (PolyPhen-2, http://genetics.bwh.harvard.edu/pph2/index.shtml) and Sorting Intolerant From Tolerant (SIFT, http://sift.jcvi.org).

### Molecular dynamics simulation of human RABEP2-RAB5 complex structure

The RABEP1-RAB5 complex structure, which was previously identified, was directly downloaded from the Protein Data Bank (PDB entry: 1TU3) (34). To predict three-dimensional complex structure of the RAB5-RABEP2, the human RABEP2 protein structure was obtained by AlphaFold2 (35,36) and then, using the HDOCK homology software (HDOCK server; http://hdock.phys.hust.edu.cn) (37) based on the RABEP1-RAB5 complex structure, the RABEP2-RAB5 complex structure was modeled. For structure modeling of RABEP2 containing Histidine at amino acid position 543, the Backbone-Dependent Rotamer library, which is embedded in the Pymol software (https://pymol.org/2/), was used to generate four possible rotamer confirmations of Histidine residue. Within the four possible Histidine rotamers, the most structurally dominant rotamer was chosen for study.

### Cell culture

Human microvascular endothelial cells (HMEC-1, CRL-3243, ATCC, VA) were cultured at 37°C in a 5% CO_2_ humidified incubator using MCDB-131 medium containing microvascular growth supplement (MVGS), 10mM L-Glutamine, and 5% FBS.

### *In vitro* scratch assay

Prior to the cell culture, vertical and horizontal reference lines were made on the bottom of 0.1% gelatin coated 6-well plate to obtain the same field for each image acquisition. Then, the HMECs were seeded in the plate at a concentration of 8×10^4^ cells/well and incubated 24 hours. To reduce endogenous human *RABEP2* mRNA expression by RNA interference, 10μM of non-specific siRNA (D-001910-10-05, Dharmacon, Inc., CO) or the human *RABEP2*-specific siRNA (M-009001-01-0005, Dharmacon, Inc., CO) was treated as recommended and the cells were maintained at 37°C in a 5% CO_2_ humidified incubator. Forty-eight hours after siRNA treatment, different AAV constructs were infected for additional 48 hours. When the infected HMECs reached over 95% confluence in 6-well plate, the wells were scratched with a 200 μL sterile pipette tip. The cells were then washed with pre-warmed PBS to remove detached cells and pre-warmed regular culture media was added. Using the reference lines as guides, the plate was placed under a phase-contrast microscope (EVOS FL, Thermo Fisher Scientific, MA) and the wound scratch was photographed at 0, 4 hours, 8 hours, and 10 hours post wounding. Wound area was calculated by manually tracing the cell-free area in captured images using the Fiji (ImageJ). The wound closure rate was expressed as the percentage of area reduction.

### Quantitative PCR (qPCR)

The mRNA levels were measured by qPCR as previously described (26,27). To quantitate mRNA levels of *RABEP2*, HMECs treated either control siRNA or si-h*RABEP2* were used. All samples were run in triplicate, and an additional assay for endogenous *GAPDH* was performed to control for input cDNA template quantity.

### Western blot analysis

To determine siRNA efficiency, protein samples were collected from HMECs treated either control siRNA or si-h*RABEP2*. In addition, to measure human RABEP2 expression levels, protein samples were collected from HMECs infected with either CMV-control (control) or CMV-h*RABEP2* (rescued) after si-*RABEP2* treatment. Briefly, HMECs were incubated in cold lysis buffer (50mM Tris-HCl [pH 7.8], 150 mM NaCl, 0.2% Triton X-100) containing protease and phosphatase inhibitor cocktail (Thermo Fisher Scientific, MA). Protein samples (30-50 μg) were electrophoresed on a 12% polyacrylamide gel and then transferred on PVDF membrane using Trans-Blot Turbo Transfer System (Bio-Rad Laboratories, CA). Membranes were incubated with primary and secondary antibodies, and the level of protein was visualized via chemiluminescence (ECL Detection Kit, Thermo Fisher Scientific, MA) The following primary antibodies were used for Western Blot analysis; rabbit anti-RABEP2 (14625-1-AP, Proteintech Group, Inc, IL) and mouse anti-GAPDH (sc-32233, Santa Cruz Biotechnology, Inc., TX). HRP-conjugated anti-mouse and anti-rabbit (Santa Cruz Biotechnology, Inc., TX) secondary antibodies were used to detect proteins.

### Statistics

Statistical analyses were performed with Prism (GraphPad Software, La Jolla, CA). Significant differences between data sets were determined using either a 2-tailed Student’s *t* test when comparing 2 groups or one-way ANOVA followed by Tukey’s test for multiple comparisons. Data are represented as the mean ± SD. *P* < 0.05 was considered significant. Table S1 provides statistical analyses for all data shown in Figures.

## Results

### Absence of *Rabep2* modulates both cerebral collateral vessel anatomy and ischemic infarct volume

To investigate whether *Rabep2* is the major modulator of both pial collateral vessel density and infarct volume, we generated *Rabep2* KO mice using CRISPR/Cas9 technology. We first examined the effect of the loss of *Rabep2* on collateral vessel connections between the anterior cerebral artery (ACA) and the middle cerebral artery (MCA), and infarct volume after pMCAO. In concordance with a previous report (32), collateral vessel connections were dramatically reduced with loss of the *Rabep2* (Fig 1A-G) leading to a concomitant increase in infarct size (Fig 1H-K) compared to both *Rabep2* WT and Het animals. This is consistent with a collateral vessel-dependent effect for *Rabep2* in modulating ischemic stroke infarction. Our data corroborate recent studies showing that *Rabep2* modulates infarct volume via the collateral circulation (Fig 1G and K).

**Figure 1.**
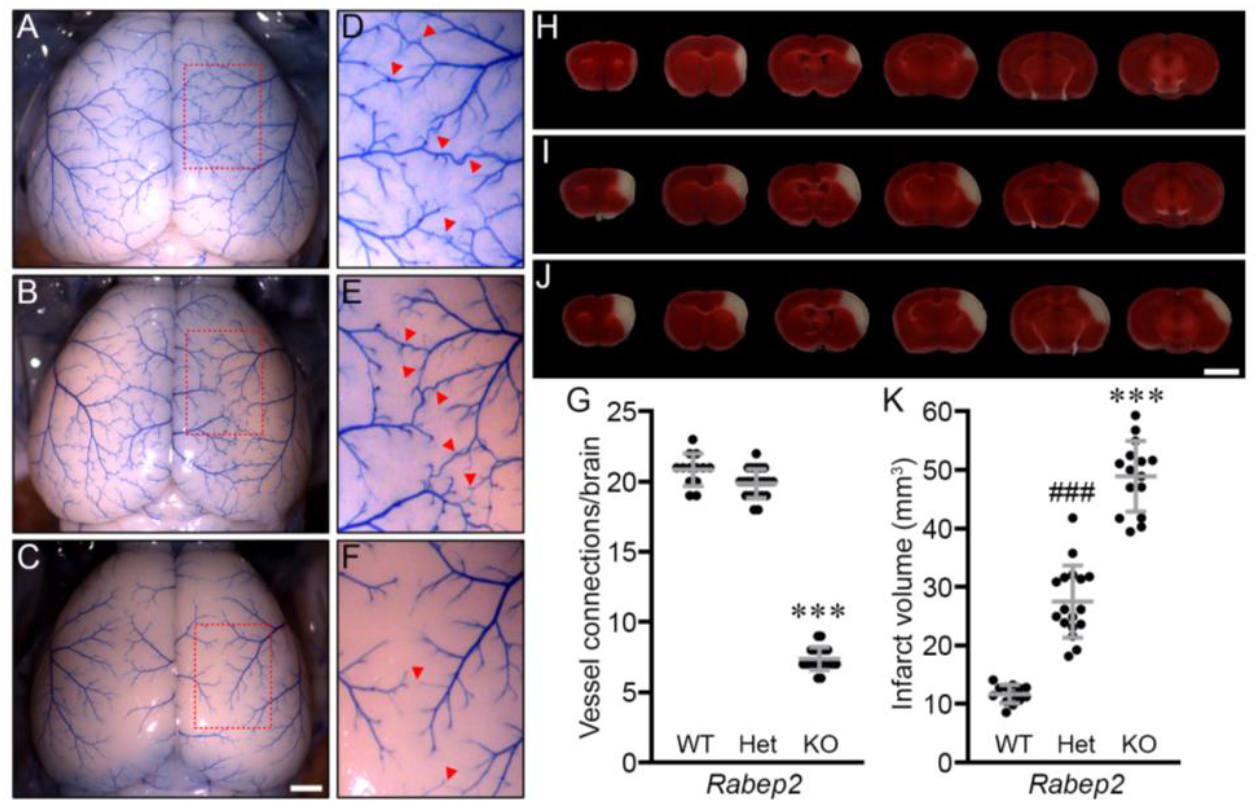
Collateral vessel density and infarct volume for *Rabep2* KO mice confirm a role for the gene in both phenotypes. **(A-C)** Representative images are shown for *Rabep2*, wild-type (WT) (A), heterozygous KO (Het) (B), and homozygous KO (KO) (C) strains. Scale bar: 1 mm. **(D-F)** D, E, and F are three times magnified from A, B, and C, respectively, and red arrowheads indicated collateral vessel connections between the ACA and MCA. **(G)** The scatter plots show the number of collateral vessel connections between the ACA and MCA in the brain for each animal. The total number of animals for *Rabep2* WT, Het, and KO were 13, 23, and 21 mice, respectively. Data represent the mean ± SD and statistical significance was determined by one-way ANOVA followed by Tukey’s multiple comparison test (*** *p* < 0.001 *vs. Rabep2* WT and Het). **(H-J)** Serial brain sections (1 mm) for each genotype of *Rabep2* WT (H), Het (I), and KO (J) are shown. The infarct appears as white tissue after 2% TTC staining. Scale bar: 5 mm. **(K)** The scatter plots show the infarct volume 24 hours after pMCAO for each animal. The total number of animals for *Rabep2* WT, Het, and KO were 13, 17, and 15 mice, respectively. Data represent the mean ± SD and statistical significance was determined by one-way ANOVA followed by Tukey’s multiple comparison test (*** *p* < 0.001 *vs. Rabep2* WT and Het; ### *p* < 0.001 *vs. Rabep2* WT and KO).

### Expression of RABEP2 rescues cerebral collateral vessel connections as well as infarct volume phenotypes

To further validate the role of *Rabep2* in cerebral infarction, we sought an approach that could restore the functional effects of the gene. We chose an “add-back” gene rescue strategy, where, in the gene knockout mice, we could express exogenous *Rabep2* delivered via recombinant AAV. To determine the efficiency of the AAV delivery system, we expressed mouse RABEP2 and GFP under the control of the CMV promoter yielding ubiquitous expression. We first evaluated RABEP2 expression *in vivo* using AAV9-CMV-mouse *Rabep2* (CMV-m*Rabep2*) that was injected directly into the right hemisphere of the brain of P1 *Rabep2* KO mice (Fig 2A). Since the AAV construct expresses both RABEP2 and GFP, GFP serve as a proxy for RABEP2 expression. Within days of injection, GFP is stably and efficiently expressed in the entire mouse brain (Fig 2B). Stronger GFP signal was observed in the right hemisphere (*Ipsilateral*) compared to the left hemisphere (*Contralateral*), providing an internal control. This expression pattern demonstrates that recombinant AAV provides an ideal platform for gene rescue in our mouse model of ischemic stroke.

**Figure 2.**
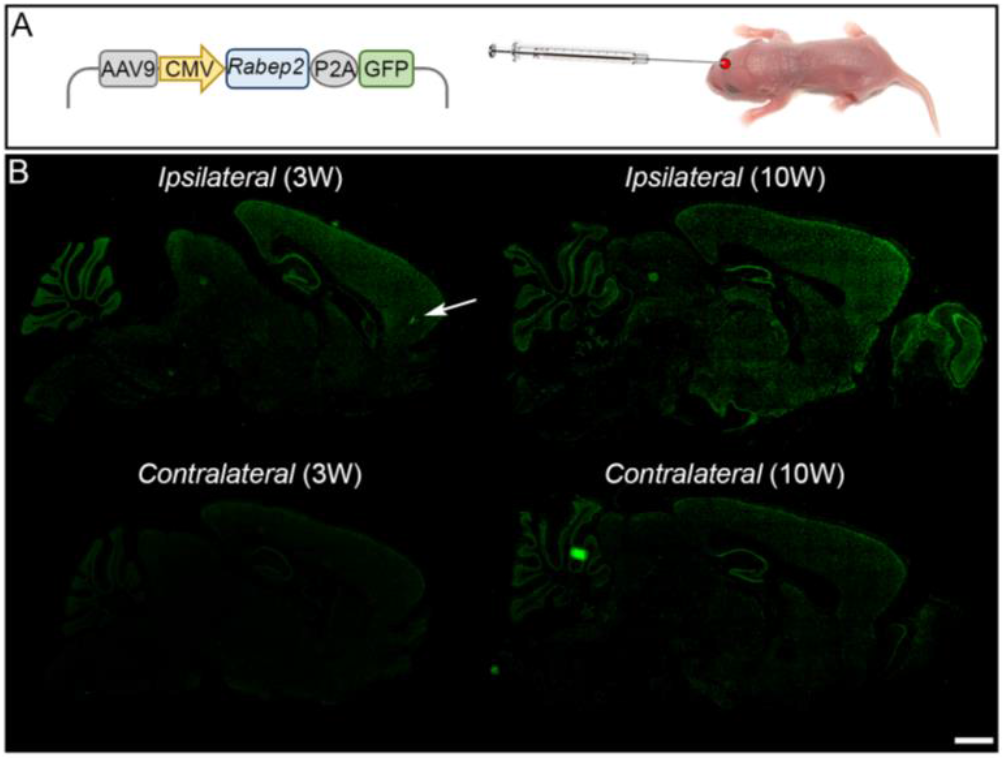
Genetic cargo of a single *in vivo* injection of recombinant AAV is widely expressed for many weeks in the mouse brain. **(A)** Schematic map of AAV containing mouse *Rabep2* and GFP driven by a CMV (ubiquitous) promoter. The recombinant AAV was injected into a P1 mouse brain (right hemisphere). **(B)** Immunostaining for recombinant GFP expressed upon AAV injection, 3 and 10 weeks after AAV9-CMV-m*Rabep2* injections. The white arrow indicates where the virus was injected into P1 mouse brain. Scale bar: 2 mm.

We first determined whether either the collateral vessel or infarct volume phenotypes were affected by injection of this AAV vector platform. There was no difference in the collateral vessel number of *Rabep2* KO mice injected with AAV empty vector containing GFP (CMV-control) compared to the *Contralateral* hemisphere (3 weeks and 10 weeks) (Fig S1A). Furthermore, CMV-control injected *Rabep2* WT, Het, and KO mice showed a similar number of collateral vessel connections as *Rabep2* WT, Het, and KO mice with no virus (Table S1). We then examined the effect of the AAV injection on ischemic infarct volume in the permanent occlusion model. Twelve weeks post injection with the CMV-control, the infarct volumes in *Rabep2* WT, Het, and KO animals, were consistent with the infarct volumes in *Rabep2* WT, Het, and KO mice with no virus (Fig1 K and Fig S1B). Thus, the injection of AAV and expression of GFP have no effect on these phenotypes.

Using this platform, we attempted to rescue the phenotype in *Rabep2* KO mice with AAV vectors containing mouse *Rabep2* under the control of a CMV promoter (Fig 3A). The brains of *Rabep2* KO mice injected with CMV-m*Rabep2* showed a time-dependent increase in collateral vessel connections compared to the *Contralateral* hemisphere as well as when compared to *Rabep2* KO mice injected with CMV-control (Fig 3B, C, and E). *Rabep2* KO mice injected with CMV-m*Rabep2* exhibited a dramatic reduction of infarct volume after pMCAO compared to *Rabep2* KO mice injected with CMV-control (Fig 3F, G, and I), consistent with the role of collateral vasculature in promoting reperfusion of the ischemic territory thereby reducing infarct volume.

**Figure 3.**
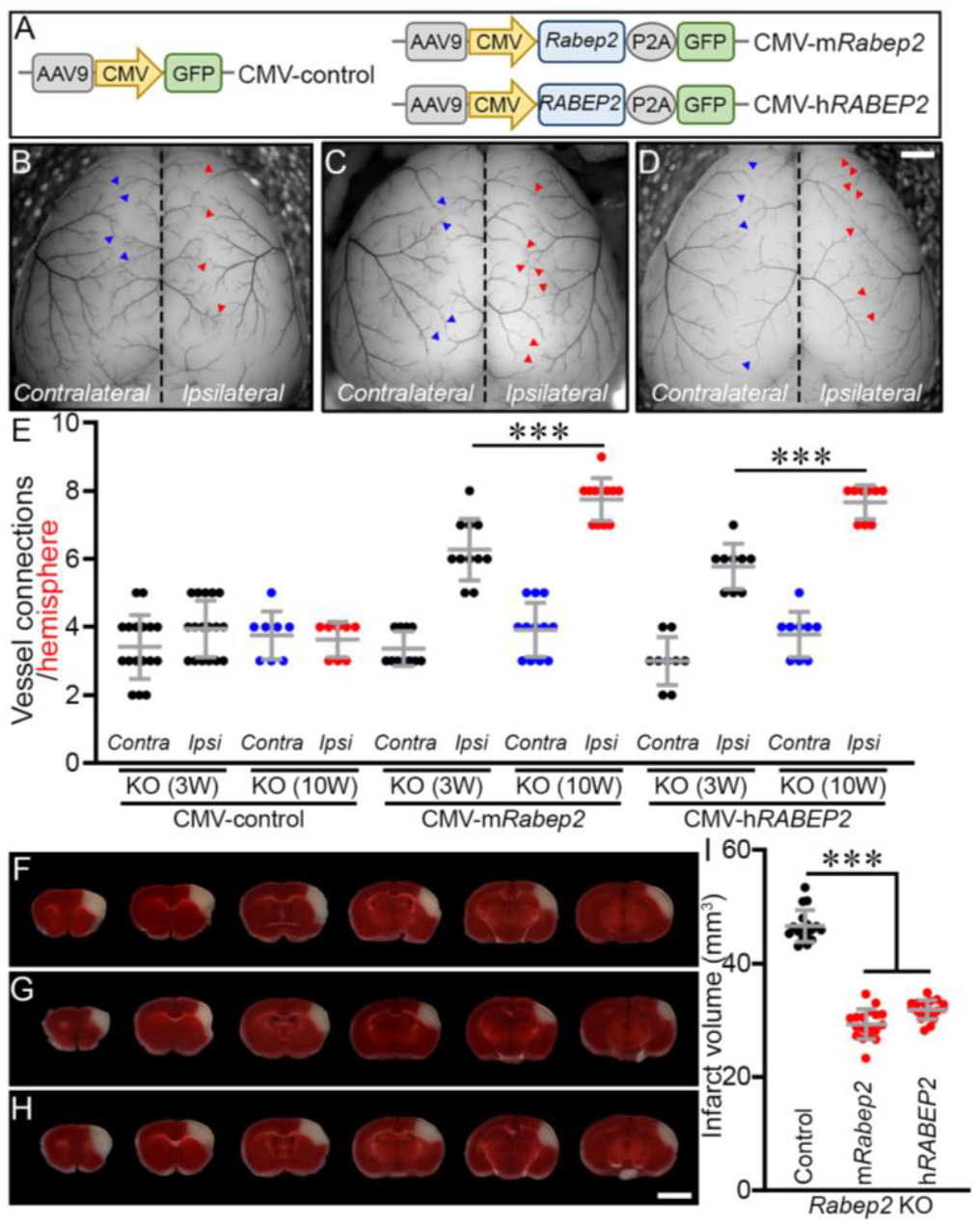
Exogenous mouse RABEP2 rescues the collateral vessel connection phenotype as well as the infarct volume phenotype of *Rabep2* KO mice. **(A)** Schematic map of AAV constructs. **(B-D)** Representative images of *Rabep2* KO mice injected with either CMV-control (B), CMV-m*Rabep2* (C), or CMV-h*RABEP2* (D). Blue and red arrowheads indicate collateral vessel connections either on the *Contralateral* (no virus) or the *Ipsilateral* (with viruses containing only GFP (B), mouse *Rabep2* (C), and human *RABEP2* (D), respectively) hemisphere. Scale bar: 1 mm. **(E)** The scatter plots show the number of collateral vessel connections between the ACA and MCA of each hemisphere, 3 and 10 weeks after AAV injections. The total number of animals for *Rabep2* KO either 3 or 10 weeks after CMV-control, *Rabep2* KO either 3 or 10 weeks after CMV-m*Rabep2*, and *Rabep2* KO either 3 or 10 weeks after CMV-h*RABEP2* are 17, 8, 11, 12, 9, and 9 animals, respectively. Data represent the mean ± SD and statistical significance was determined by one-way ANOVA followed by Tukey’s multiple comparison test (*** *p* < 0.001). **(F-H)** Serial brain sections (1 mm) for *Rabep2* KO mice injected with either CMV-control (F), CMV-m*Rabep2* (G), or CMV-h*RABEP2* (H). Scale bar: 5 mm. **(I)** The scatter plots show the infarct volume for *Rabep2* KO mice injected with CMV-control, CMV-m*Rabep2*, and CMV-h*RABEP2*; 17, 18, and 18 animals, respectively. Data represent the mean ± SD and statistical significance was determined by one-way ANOVA followed by Tukey’s multiple comparison test (*** *p* < 0.001).

Although the collateral vessel network is thought to be set during embryonic collaterogenesis (32), these data show that new collateral vessels also develop in early postnatal life (Fig 3E). We note however, that there was no increase in collateral vessel connections on the *Contralateral* hemisphere even 10 weeks after virus injection (Fig 3C and E), despite what appears to be relatively strong expression of *Rabep2* through the entire brain (Fig 2B). This suggests a narrow threshold for either gene dose or timing, or both, for collateral vessel formation after birth.

### Human RABEP2 rescues the collateral vessel and infarct volume phenotypes of *Rabep2* KO mice

We next determined whether human RABEP2 could rescue the KO phenotypes as we observed with mouse *Rabep2* (Fig 3E and I). We expressed the human *RABEP2* gene (h*RABEP2*) in the same AAV platform (CMV-h*RABEP2*) (Fig 3A), and we determined the effect of injection of h*RABEP2* in *Rabep2* KO mice. Expression of human RABEP2 increased collateral vessel number (Fig 3D and E) and reduced infarct volume of *Rabep2* KO mice upon ischemic stroke (Fig 3H and I). Within 3 weeks, the rescued KO mice had established new collateral vessel connections, and by 10 weeks the number of these connections increased further. At 12 weeks of age, upon stroke induction, the resulting infarct volume was also reduced in the rescued *Rabep2* KO mice. Thus, the human *RABEP2* gene rescues the *Rabep2* KO mouse phenotypes. Furthermore, both human *RABEP2* and mouse *Rabep2* genes rescue collateral vessel and infarct volume phenotypes to similar levels (Fig 3). This strongly supports the hypothesis that the *Rabep2* gene shares the same function in both humans and mice.

### Prediction of functional consequences of human RABEP2 coding variants

Since we confirmed that human RABEP2 is fully functional in the mouse, we investigated whether coding sequence variants of the human *RABEP2* impact ischemic infarction. If coding variants in this gene alter infarct volume *in vivo*, this gene gains further “human genetics” support as a therapeutic target. Using the full list of RABEP2 coding SNPs found in the Genome Aggregation Database (gnomAD (v2.1.1)), we first ordered all coding SNP alleles based on overall population allele frequency. Then, we examined the predicted functional consequences of these SNPs by two independent *in silico* algorithms, SIFT and PolyPhen-2. Using these algorithms, we prioritized our initial functional analysis to three coding SNP variants that are strongly predicted to be damaging (Table 1; highlighted in yellow).

**Table 1.** *In silico*-predicted functional consequences of coding variants of human RABEP2. The list is based on human population data obtained from the Genome Aggregation Database (gnomAD) and functional effects of coding SNPs were predicted by two independent *in silico* algorithms, SIFT and PolyPhen-2. Red highlight indicates a strong prediction of functional damage. Orange indicates possibly damaging. A yellow highlighted indicates the four variants chosen for further study.

| Chr | Position | rsID | ref | alt | Coding SNP | Allele |  |  | *In silico* prediction |  |
| --- | --- | --- | --- | --- | --- | --- | --- | --- | --- | --- |
|  |  |  |  |  |  | count | number | frequency | SIFT | PolyPhen |
| 16 | 28936427 | rs115529621 | C | A | A20S | 3562 | 167926 | 0.021211724 | 0.16 | 0 |
|  | 28916802 | rs200118396 | C | A | R508S | 239 | 271860 | 0.000879129 | 0 | 0.968 |
|  | 28922259 | rs201270750 | T | A | T347S | 137 | 278198 | 0.000492455 | 0.27 | 0.145 |
|  | 28922244 | rs202177661 | C | A | V352L | 121 | 278060 | 0.000435158 | 0.36 | 0.015 |
|  | 28917045 | rs201409528 | C | T | V491M | 111 | 280082 | 0.000396313 | 0.09 | 0.207 |
|  | 28917367 | rs559738470 | G | C | Q466E | 107 | 219272 | 0.000487978 | 1 | 0.006 |
|  | 28925840 | rs769480150 | G | A | S204L | 91 | 275124 | 0.00033076 | 0.01 | 0.978 |
|  | 28917045 | rs201409528 | C | A | V491L | 80 | 280082 | 0.000285631 | 0.42 | 0.026 |
|  | 28919991 | rs192852294 | G | A | S395L | 80 | 278986 | 0.000286753 | 0.84 | 0 |
|  | 28916806 | rs766728154 | T | A | Q507L | 57 | 272456 | 0.000209208 | 0 | 0.127 |
|  | 28931252 | rs199904424 | C | T | S96N | 49 | 272106 | 0.000180077 | 0.14 | 0.988 |
|  | 28931168 | rs758567963 | C | T | R124H | 42 | 278940 | 0.00015057 | 0.29 | 0.019 |
|  | 28917048 | rs184144701 | G | A | R490W | 34 | 280036 | 0.000121413 | 0 | 0.995 |
|  | 28922430 | rs779592285 | C | T | R322Q | 32 | 280052 | 0.000114264 | 0.33 | 0.36 |
|  | 28925870 | rs372536339 | G | A | T194M | 26 | 275034 | 9.45338E-05 | 0.16 | 0.513 |
|  | 28925859 | rs764221819 | G | A | P198S | 24 | 244096 | 9.8322E-05 | 0.28 | 0.003 |
|  | 28917442 | rs750752217 | G | A | R441C | 21 | 206462 | 0.000101714 | 0 | 0.993 |
|  | 28925703 | rs372007568 | G | A | P250S | 21 | 277996 | 7.55407E-05 | 0.07 | 0.063 |
|  | 28925904 | rs550518363 | G | A | R183C | 21 | 273966 | 7.66518E-05 | 0.02 | 0.549 |
|  | 28925694 | rs200278634 | G | A | R253C | 20 | 278590 | 7.17901E-05 | 0.03 | 0.828 |
|  | 28916281 | rs376299848 | C | A | D565Y | 19 | 279334 | 6.80189E-05 | 0 | 0.978 |
|  | 28926097 | rs199981658 | C | G | E147Q | 19 | 249182 | 7.62495E-05 | 0.01 | 0.945 |
|  | 28916280 | rs759288035 | T | A | D565V | 17 | 279320 | 6.08621E-05 | 0 | 0.93 |
|  | 28917492 | rs369261936 | G | A | T424M | 17 | 234784 | 7.2407E-05 | 0.21 | 0.104 |
|  | 28916346 | rs527458355 | C | T | R543H | 16 | 273404 | 5.85215E-05 | 0 | 0.999 |
|  | 28922226 | rs748834926 | G | A | R358W | 16 | 277234 | 5.7713E-05 | 0 | 0.948 |
|  | 28925850 | rs375415976 | G | A | R201W | 16 | 275346 | 5.81087E-05 | 0.03 | 0.006 |

We sought to identify additional potential candidate coding variants by focusing on the cellular role of RABEP2 which regulates membrane trafficking in the early endocytic pathway with RAB5 (38). A previous study has reported that human RABEP1 interacts with RAB5 and this interaction complex is disrupted by the mutation of any one of the interaction residues (D820, Q826, and Q829) in RABEP1 (34) (Fig S2A). To further identify potential interaction residue(s) between RABEP1 and RAB5, we performed structural analysis using the RABEP1-RAB5 complex structure, and then identified four amino acid residues (V817, L824, E833, and Q837) in RABEP1 (Fig S2A). These four amino acid residues in RABEP1 have also been suggested as interaction residues with RAB5 (34). Since human RABEP1 and 2 share a high protein sequence identity and both interact with RAB5 (38), we aligned the C-terminal RAB5 binding motifs of both RABEP1 and 2 that share over 80% amino acid similarity (Fig S2B). We found that all seven interaction residues (experimentally confirmed (red) and structurally predicted (blue)) are conserved between RABEP1 and RABEP2 (Fig S2A and B). To analyze the RABEP2-RAB5 complex structure, we performed molecular dynamics simulation. We first obtained human RABEP2 protein structure from AlphaFold 2, an artificial intelligence-based protein structure prediction program (35,36). Then, using HDOCK software to predict the binding complexes between two molecules (37), we modeled the RABEP2-RAB5 complex structure. Based on these structural and *in silico* analyses, we identified additional coding SNP variants from the human population database that lie at or near these known or predicted contact points. A coding SNP variant at amino acid position 543, substituting Histidine (H) for Arginine (R) (Table 1; highlighted in yellow), is predicted to be damaging due to disruption of critical ionic interactions (Fig 4A-C). This variant is also predicted to be damaging by the *in silico* prediction algorithms. This variant became our fourth variant for study.

**Figure 4.**
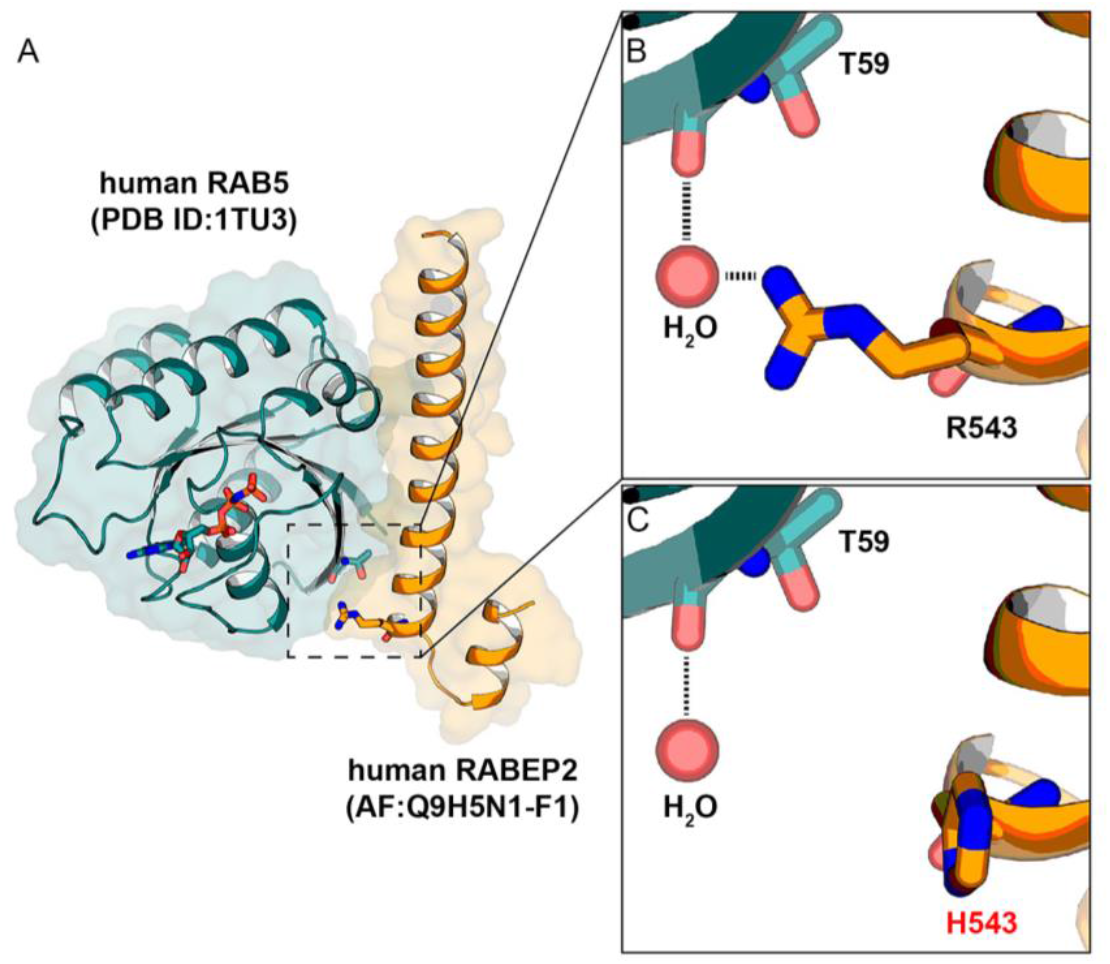
Protein-protein docking between human RABEP2 and human RAB5. **(A)** Homologous modeling of the interaction of human RABEP2 with human RAB5 showing the secondary structure using HDOCK homology software. **(B and C)** Closed-up views of the interacting interfaces of human RAB5 with human RABEP2 (R543) (B) and human RAB5 with human RABEP2 coding variant (H543) (C). The black dotted-lines indicate water-mediated hydrogen bonds. A water molecule is shown as a red sphere.

To investigate the potential effects of the four human coding SNP variants of RABEP2, we generated recombinant AAV constructs carrying these putative loss-of-function sequence variants along with the consensus coding sequence as a control. We validated the sequence of each variant by the Sanger sequencing (Fig S3A-D).

### Human RABEP2 coding variants reduce human endothelial cell migration

To investigate the role of RABEP2 on endothelial cell biology *in vitro*, we examined the effects of the gene on endothelial cell migration (Fig 3A and Fig S3). We first determined the extent of protein loss by acute knockdown of h*RABEP2* gene expression in human microvascular endothelial cells (HMECs) using h*RABEP2*-specific siRNA (si-h*RABEP2*). The efficiency of siRNA delivery in HMECs was evaluated for up to 72 hours. h*RABEP2* mRNA transcript levels in HMECs showed a significant and gradual reduction after transfection of si-h*RABEP2* compared to that observed after transfection with nonspecific siRNA (control siRNA) (Fig S4A). RABEP2 protein levels concomitantly decreased (Fig S4B). At the 48-hour time point after siRNA treatment, we confirmed that we could restore RABEP2 protein levels for an additional 48 hours after infection with CMV-h*RABEP2*. Levels increased approximately 12-fold compared to control (CMV-control) (Fig S4C).

To determine whether human RABEP2 coding variants would affect endothelial cell migration, we performed the *in vitro* scratch assay (i.e., cell migration) after knockdown of h*RABEP2* expression by si-h*RABEP2* in HMECs, expressing different AAV constructs (Fig 5). As shown in Figure 5A, HMECs expressing CMV-control after treatment of si-h*RABEP2* (Loss-of-Function; LoF) showed dramatically reduced cell migration as compared to HMECs treated with control siRNA and infected with the CMV-control. By contrast, the cell migration was rescued when exogenous RABEP2 was delivered (Fig 5A and B) indicating that RABEP2 is able to influence endothelial cell migration. The cell migration in HMECs expressing RABEP2 with the variant R543H was drastically reduced (Fig 5A and B), comparable to the migration of the LoF si-h*RABEP2*. We also performed the cell migration assay using the three other human RABEP2 coding variants (R508S, S204L, and R490W). All variants showed similar levels of cell migration rate to R543H (Fig 5B and C), suggesting these variants also represent hypomorphic or loss-of-function alleles that segregate in the human population. These data identify another important cellular function of RABEP2 in endothelial cells.

**Figure 5.**
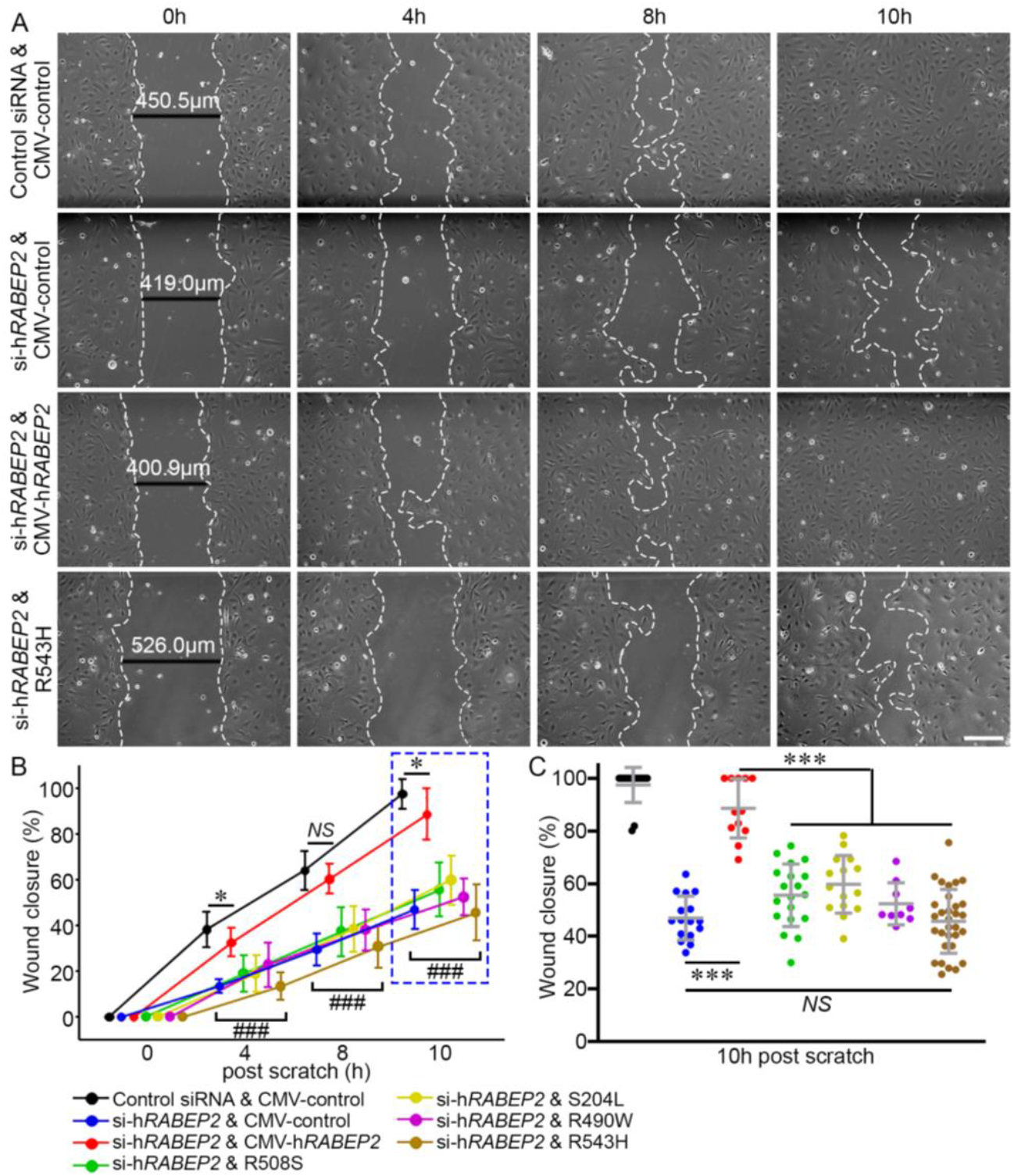
Human RABEP2 coding variants impair cell migration in an *in vitro* scratch wound assay. **(A)** Representative bright-field images display 4 different groups; Control infected with CMV-control after control siRNA treatment, Loss-of-Function (LoF) infected with CMV-control after si-h*RABEP2* treatment, Rescued infected with CMV-h*RABEP2* after si-h*RABEP2* treatment, and R543H infected with CMV-h*RABEP2*-R543H after si-h*RABEP2* treatment. A fully confluent (100%) HMEC monolayer for each treatment was scratched with the tip of a 200 μl pipet. At different time intervals (0, 4, 8, and 10 h), the degree of migration of HMECs was imaged. **(B)** The line graph shows wound closure ratio. The wound closure rate was quantified as the percentage of area reduction at each time point. The total number of scratch experiments for Control, LoF, Rescued, R508S, S204L, R490W, and R543H were 15, 16, 12, 19, 15, 9, and 31, respectively. Data represent the mean ± SD and statistical significance was determined by one-way ANOVA followed by Tukey’s multiple comparison test (* *p* < 0.05; ### (individual group) *p* < 0.001 *vs*. control and rescued groups; *NS*, not significant). **(C)** The scatter plots show the percentage of wound closure 10 hours after scratch. Data represent the mean ± SD and statistical significance was determined by one-way ANOVA followed by Tukey’s multiple comparison test (*** *p* < 0.001).

### Human RABEP2 coding variants abolish the ability of RABEP2 to rescue the collateral vessel and infarct volume phenotypes in *Rabep2* KO mice

To investigate whether human RABEP2 coding variants could modulate clinically relevant phenotypes of collateral vessel number and ischemic stroke infarct volume, we examined these phenotypes in *Rabep2* KO mice expressing the different RABEP2 coding variants. We first measured the number of collateral vessel connection between the ACA and MCA in *Rabep2* KO mice. The number of collateral vessel connections in *Rabep2* KO mice injected with either CMV-h*RABEP2*-R508S, CMV-h*RABEP2*-S204L, or CMV-h*RABEP2*-R490W were slightly increased compared to *Rabep2* KO mice injected with CMV-control (Fig 6A-E), however, these collateral vessel connections were not as fully developed as those we observed in *Rabep2* KO mice injected with CMV-h*RABEP2* (Fig 6E). Interestingly, CMV-h*RABEP2*-R543H, which is predicted to disrupt the interaction between RABEP2 and RAB5, has no ability to rescue the collateral vessel defect (Fig 6E). The number of collateral vessel connections in *Rabep2* KO mice injected with CMV-h*RABEP2*-R543H is similar to *Rabep2* KO mice injected with CMV-control (Fig 6E). *Rabep2* KO mice injected with either CMV-h*RABEP2*-R508S, CMV-h*RABEP2*-S204L, or CMV-h*RABEP2*-R490W did show increased collateral vessel connections between 3 and 10 weeks after AAV injection (Fig S5). However, *Rabep2* KO mice injected with CMV-h*RABEP2*-R543H showed no difference in collateral vessel connections between 3 and 10 weeks after AAV injection (Fig S5). Thus, this coding SNP, R543H, of RABEP2 is effectively a null allele.

**Figure 6.**
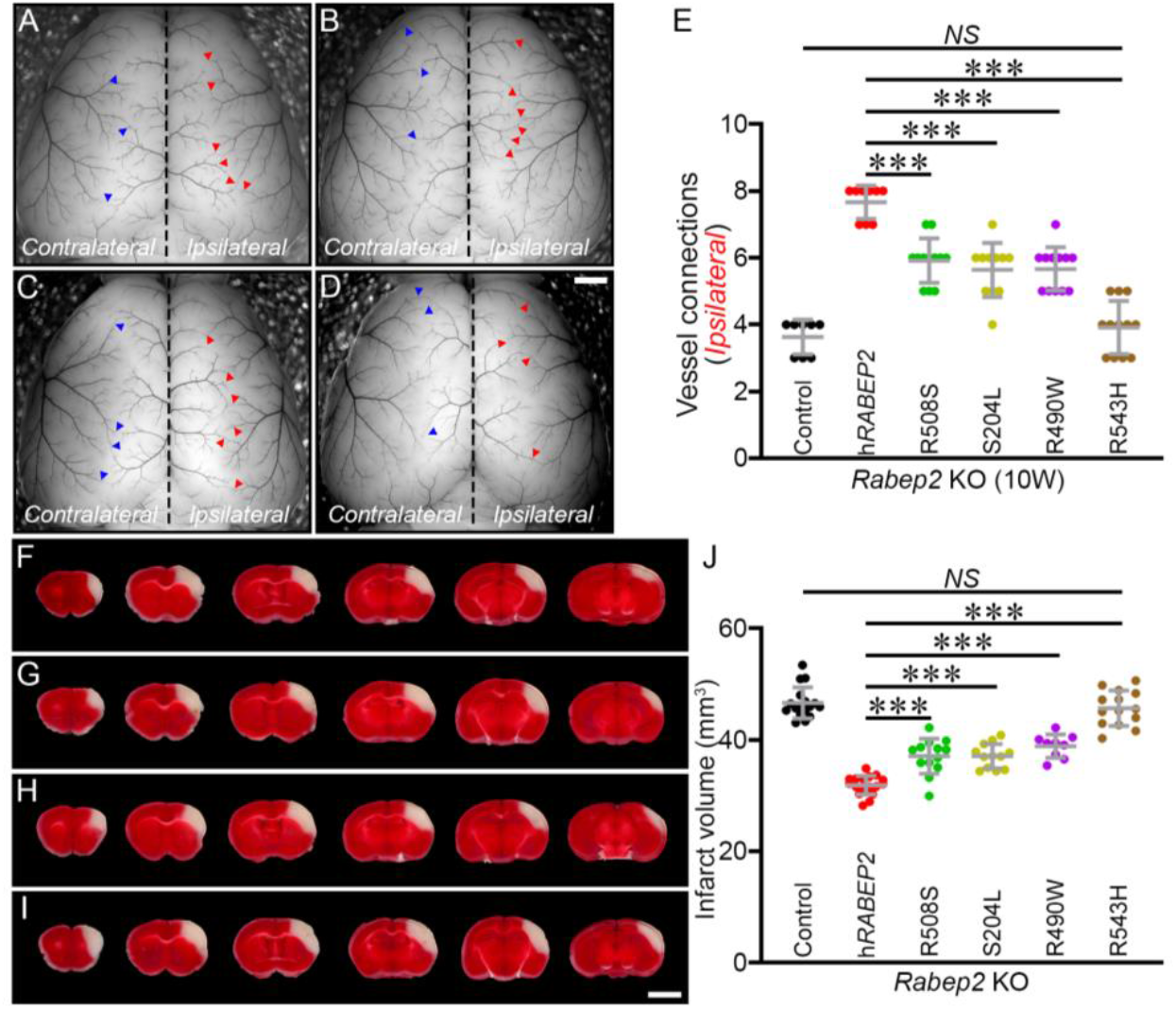
Human RABEP2 coding variants significantly impair gene rescue for both collateral vessel density and infarct volume phenotypes. **(A-D)** Representative images of *Rabep2* KO mice injected with either R508S (A), S204L (B), R490W (C), or R543H (D). Blue and red arrowheads indicate collateral vessel connections on the *Contralateral* (no virus) and on the *Ipsilateral* (with virus) hemisphere, respectively. Scale bar: 1 mm. **(E)** The scatter plots show the number of collateral vessel connections between ACA and MCA of the *Ipsilateral* hemisphere, 10 weeks after AAV injection. The total number of animals for *Rabep2* KO injected with CMV-control, CMV-h*RABEP2*, R508S, S204L, R490W, and R543H are 8, 9, 12, 11, 12, and 12 animals, respectively. Data represent the mean ± SD and statistical significance was determined by one-way ANOVA followed by Tukey’s multiple comparison test (*** *p* < 0.001; *NS*, not significant). **(F-I)** Serial brain sections (1 mm) for *Rabep2* KO mice injected with either R508S (F), S204L (G), R490W (H), or R543H (I). Scale bar: 5 mm. **(J)** The scatter plots show the infarct volume for *Rabep2* KO mice injected with CMV-control, CMV-h*RABEP2*, R508S, S204L, R490W, and R543H; 17, 18, 13, 12, 9, and 14 animals, respectively. Data represent the mean ± SD and statistical significance was determined by one-way ANOVA followed by Tukey’s multiple comparison test (** *p* < 0.01; *** *p* < 0.001; *NS*, not significant).

We next examined the human RABEP2 coding variants on ischemic stroke in the permanent occlusion model. We introduced each human RABEP2 coding variant with recombinant AAV injection into *Rabep2* KO mice and performed pMCAO to measure infarct volume for each individual coding variant (Fig 6F-I). As observed the collateral vessel phenotypes in *Rabep2* KO mice with different coding variants, the infarct volume in *Rabep2* KO mice injected with either CMV-h*RABEP2*-R508S, CMV-h*RABEP2*-S204L, or CMV-h*RABEP2*-R490W was reduced compared to *Rabep2* KO mice injected with CMV-control (Fig 6J). However, these infarct volumes were still larger than that observed in *Rabep2* KO mice injected with CMV-h*RABEP2*, suggesting that these coding variants are hypomorphic alleles (Fig 6J). The nfarct volume in *Rabep2* KO mice injected with CMV-h*RABEP2*-R543H showed no difference with *Rabep2* KO mice injected with CMV-control (Fig 6J), Together with the results from the rescue experiment for collateral vessel number, it appears that this variant is a null allele. These data show that certain human RABEP2 coding variants are either hypomorphs or complete loss-of-function alleles for both phenotypes of collateral vessel connections and infarct volume.

The overall population allele frequencies for these four variants are quite low. While R508S is found in multiple populations, the other three of these variants exclusively segregate within distinct populations. As shown in Table 2, S204L, R490W, and R543H are predominantly found in the European, East Asian, and African/African American population, respectively. These coding variants are potentially a strong determinant of stroke outcome in these populations.

**Table 2.** Population frequencies for four coding variants in human RABEP2. The list displays population-based allele frequencies of coding SNP variants. While one coding SNP (R508S) in human RABEP2 is widely distributed across populations, coding SNPs S204L, R490W, or R543H exclusively segregate either the European, East Asian, or African/African American population.

| Population | R508S |  |  | S204L |  |  | R490W |  |  | R543H |  |  |
| --- | --- | --- | --- | --- | --- | --- | --- | --- | --- | --- | --- | --- |
|  | Count | Allele Num. | Freq. | Count | Allele Num. | Freq. | Count | Allele Num. | Freq. | Count | Allele Num. | Freq. |
| Ashkenazi Jewish | 29 | 10036 | 0.00289 | 0 | 10168 | 0 | 0 | 10326 | 0 | 0 | 10268 | 0 |
| European (non-Finnish) | 136 | 124560 | 0.001092 | 7 | 124038 | 0.0000564 | 0 | 128000 | 0 | 0 | 125522 | 0 |
| European (Finnish) | 1 | 23762 | 0.0000421 | **84** | **24870** | **0.003378** | 0 | 24992 | 0 | 0 | 21320 | 0 |
| African/African American | 2 | 23348 | 0.0000857 | 0 | 23748 | 0 | 2 | 24130 | 0.0000829 | **15** | **23982** | **0.0006255** |
| Latino/Admixed American | 22 | 34648 | 0.000635 | 0 | 35250 | 0 | 0 | 35346 | 0 | 0 | 35236 | 0 |
| East Asian | 1 | 19204 | 0.0000521 | 0 | 19486 | 0 | **31** | **19514** | **0.001589** | 0 | 19490 | 0 |
| South Asian | 34 | 29398 | 0.001157 | 0 | 30506 | 0 | 1 | 30600 | 0.0000328 | 1 | 30534 | 0.0000327 |
| Other | 14 | 6904 | 0.002028 | 0 | 7058 | 0 | 0 | 7128 | 0 | 0 | 7052 | 0 |
| XX | 91 | 123556 | 0.0007365 | 44 | 124658 | 0.000353 | 17 | 127538 | 0.0001333 | 9 | 123862 | 0.0000727 |
| XY | 148 | 148304 | 0.000998 | 47 | 150466 | 0.0003124 | 17 | 152498 | 0.0001115 | 7 | 149542 | 0.0000468 |
| Total | 239 | 271860 | 0.0008791 | 91 | 275124 | 0.0003308 | 34 | 280036 | 0.0001214 | 16 | 273404 | 0.0000585 |

## Discussion

Although mouse models of ischemic stroke have been widely used to gain insight into pathobiological mechanisms, translation of these findings to the clinic have been largely disappointing (39-41). As an alternative strategy, identification of drug targets has more recently focused on candidate gene identification through human genetics (42). Therapeutic targets identified due to their genetic variants that predict risk or disease outcome are twice as likely to progress through the phases of clinical development as those identified through other experimental approaches (43,44).

In this study, we focused on investigating and validating a potential candidate gene for ischemic stroke, *Rabep2*, by examining the effects of human sequence variants on clinically relevant phenotypes. The mouse *Rabep2* gene was identified using a forward genetic mapping approach in mouse inbred strains (23,24,26). And a single locus containing the mouse *Rabep2* gene was identified as modulating infarct volume through a collateral vessel-dependent pathway (32). Yet to date, there is no evidence that human *RABEP2* plays a role in either ischemic stroke or stroke outcomes. Instead, GWAS studies have shown that SNPs near or within the *RABEP2* gene are associated with body mass, obesity, type 2 diabetes, and neurological phenotypes (e.g., Parkinson’s disease, intelligence, cortical surface size) (45-51).

We have developed an *in vivo* evaluation platform to investigate the functional effects of human sequence variations in *RABEP2*. This evaluation platform is based on an “add-back” gene rescue strategy and an *in vivo* rescue platform. We express the missing gene in knockout mice using recombinant AAV. Employing this *in vivo* evaluation platform, we show that we can rescue the *Rabep2* KO phenotype. Significantly, we also show that the human *RABEP2* rescues the mouse *Rabep2* KO ischemic stroke phenotypes. This cross-species complementation allows us to experimentally investigate the functional effects of sequence variation in the human gene.

We have identified 4 different coding variants in human RABEP2 that were predicted to be damaging by *in silico* prediction algorithms. One of these was further identified as potentially damaging by structural analyses. Although relatively rare, three of these are population specific variants. Coding variant R543H, which lies near a critical interaction residue for RAB5, appears to be a complete loss-of-function allele. This allele segregates exclusively in the African/African American population.

We examined the effects of these variants on an endothelial cell biological phenotype (scratch wound migration) and *in vivo*, in the rescue of the phenotypes of collateral vessel number and infarct volume after pMCAO. When injected into the *Rabep2* KO mice, we showed that each of these four variants are at best hypomorphic alleles, with one variant (R543H) appearing as a null allele for all three phenotypes. *In silico* analyses have further identified other coding variants in *RABEP2* that are predicted to be damaging. Thus, *RABEP2* appears to be rich in potentially damaging coding variations that may influence human stroke outcomes. By cross-species validation, we show that variation of the human *RABEP2* gene might play a role in human ischemic stroke by modulating infarct volume through effects of the vascular circulation. We note, however, that in the *Rabep2* KO mice, the two ischemic stroke phenotypes, collateral vessel density and infarct volume, are differentially rescued by different *RABEP2* coding variants, suggesting a more complex role for this gene in ischemic stroke.

This study again demonstrates the importance *Rabep2* on ischemic stroke outcome. We have also verified RABEP2 as a new therapeutic target candidate in human ischemic stroke although further studies are required to determine the mechanism of action of RABEP2 in ischemic stroke. Additionally, we show that an *in vivo* evaluation platform in mice using AAV gene replacement is an effective and efficient method for screening the functional effects of gene variants. Using this system, we can query the functional consequences coding variants of almost any human gene relevant to ischemic stroke in the mouse. This will expand the number of physiologically relevant target candidates with the goal of identifying therapeutics for human ischemic stroke.

## Author contributions

HKL and DAM designed research. HKL performed research. DHK performed structure modeling. HKL and DAM analyzed the data. HKL and DAM wrote the manuscript.

## Description of supplemental data

Supplemental data include five figures and one table.

## Acknowledgments

The authors thank Mr. Christian R. Benavides for animal husbandry. This study was supported by grants from NIH grant 5R01NS100866 (to DAM), the Foundation Leducq Transatlantic Network Of Excellence in Neurovascular Disease 17 CVD 03 (to DAM), and American Heart Association Career Development Award 938553 (to HKL).

## Data and code availability

This study did not generate datasets.

## References

1. Bogousslavsky J., Aarli J., Kimura J. (2003). Stroke: time for a global campaign? Cerebrovasc Dis. 16, 111–113.

2. Johnson W., Onuma O., Owolabi M., Sachdev S. (2016). Stroke: a global response is needed. Bull World Health Organ. 94, 634–634A.

3. Krishnamurthi R.V., Feigin V.L., Forouzanfar M.H., Mensah G.A., Connor M., Bennett D.A., Moran A.E., Sacco R.L., Anderson L.M., Truelsen T., et al. (2013). Global and regional burden of first-ever ischaemic and haemorrhagic stroke during 1990-2010: findings from the Global Burden of Disease Study 2010. Lancet Glob Health. 1, e259–81.

4. Kim J., Thayabaranathan T., Donnan G.A., Howard G., Howard V.J., Rothwell P.M., Feigin V., Norrving B., Owolabi M., Pandian J., et al. (2020). Global Stroke Statistics 2019. Int J Stroke. 15, 819–838.

5. Mozaffarian D., Benjamin E.J., Go A.S., Arnett D.K., Blaha M.J., Cushman M., Das S.R., de Ferranti S., Després J.-P., Fullerton H.J., et al. (2016). Heart Disease and Stroke Statistics-2016 Update: A Report From the American Heart Association. Circulation. 133, e38–360.

6. Roger V.L., Go A.S., Lloyd-Jones D.M., Benjamin E.J., Berry J.D., Borden W.B., Bravata D.M., Dai S., Ford E.S., Fox C.S., et al. (2012). Heart disease and stroke statistics-2012 update: a report from the American Heart Association. Circulation. 125, e2–220.

7. Benjamin E.J., Muntner P., Alonso A., Bittencourt M.S., Callaway C.W., Carson A.P., Chamberlain A.M., Chang A.R., Cheng S., Das S.R., et al. (2019). Heart disease and stroke statistics-2019 update: a report from the American heart association. Circulation. 139, e56–528.

8. Virani S.S., Alonso A., Benjamin E.J., Bittencourt M.S., Callaway C.W., Carson A.P., Chamberlain A.M., Chang A.R., Cheng S., Delling F.N., et al. (2020). Heart Disease and Stroke Statistics-2020 Update: A Report From the American Heart Association. Circulation. 141, e139–596.

9. Goyal M., Demchuk A.M., Menon B.K., Eesa M., Rempel J.L., Thornton J., Roy D., Jovin T.G., Willinsky R.A., Sapkota B.L., et al. (2015). Randomized assessment of rapid endovascular treatment of ischemic stroke. N. Engl. J. Med. 372, 1019–1030.

10. Kim Y.D., Nam H.S., Kim S.H., Kim E.Y., Song D., Kwon I., Yang S.-H., Lee K., Yoo J., Lee H.S., et al. (2015). Time-dependent Thrombus resolution after tissue-type plasminogen activator in patients with stroke and mice. Stroke. 46, 1877–1882.

11. Muchada M., Rodriguez-Luna D., Pagola J., Flores A., Sanjuan E., Meler P., Boned S., Alvarez-Sabin J., Ribo M., Molina C.A., et al. (2014). Impact of time to treatment on tissue-type plasminogen activator-induced recanalization in acute ischemic stroke. Stroke. 45, 2734–2738.

12. Emberson J., Lees K.R., Lyden P., Blackwell L., Albers G., Bluhmki E., Brott T., Cohen G., Davis S., Donnan G., et al. (2014). Effect of treatment delay, age, and stroke severity on the effects of intravenous thrombolysis with alteplase for acute ischemic stroke: a meta-analysis of individual patient data from randomized trials. Lancet. 384, 1929–1935.

13. Albers G.W., Goldstein L.B., Hess D.C., Wechsler L.R., Furie K.L., Gorelick P.B., Hurn P., Liebeskind D.S., Nogueira R.G., Saver J.L. (2011). STAIR VII Consortium. Stroke Treatment Academic Industry Roundtable (STAIR) recommendations for maximizing the use of intravenous thrombolytics and expanding treatment options with intra-arterial and neuroprotective therapies. Stroke. 42, 2645–2650.

14. O’Donnell M.J., Xavier D., Liu L., Zhang H., Chin S.L., Rao-Melacini P., Rangarajan S., Islam S., Pais P., McQueen M.J., et al. (2010). Risk factors for ischemic and intracerebral hemorrhagic stroke in 22 countries (the interstroke study): A case-control study. Lancet. 376, 112–123.

15. Go A.S., Mozaffarian D., Roger V.L., Benjamin E.J., Berry J.D., Borden W.B., Bravata D.M., Dai S., Ford E.S., Fox C.S., et al. (2013). Heart disease and stroke statistics-2013 update: A report from the American heart association. Circulation. 127, e6–245.

16. Elkind M.S.V. (2007). Why now? Moving from stroke risk factors to stroke triggers. Curr Opin Neurol. 20, 51–57.

17. Boehme A.K., Escnwa C., Elkind M.S.V. (2017). Stroke Risk Factors, Genetics, and Prevention. Circ. Res. 120, 472–495.

18. Gretarsdottir S., Thorleifsson G., Reynisdottir S.T., Manolescu A., Jonsdottir S., Jonsdottir T., Gudmundsdottir T., Bjarnadottir S.M., Einarsson O.B., Gudjonsdottir H.M., et al. (2003). The gene encoding phosphodiesterase 4D confers risk of ischemic stroke. Nat. Genet. 35, 131–138.

19. Helgadottir A., Manolescu A., Thorleifsson G., Gretarsdottir S., Jonsdottir H., Thorsteinsdottir U., Samani N.J., Gudmundsson G., Grant S.F.A., Thorgeirsson G., et al. (2004). The gene encoding 5-lipoxygenase activating protein confers risk of myocardial infarction and stroke. Nat. Genet. 36, 233–239.

20. Matarín M., Brown W.M., Scholz S., Simón-Sánchez J., Fung H.-C., Hernandez D., Gibbs J.R., De Vrieze F.W., Crews C., Britton A., et al. (2007). A genome-wide genotyping study in patients with ischemic stroke: initial analysis and data release. Lancet Neurol. 6, 414–420.

21. Matarin M., Brown W.M., Singleton A., Hardy J.A., Meschia J.F., ISGS investigators. (2008). Whole genome analyses suggest ischemic stroke and heart disease share an association with polymorphisms on chromosome 9p21. Stroke. 39, 1586–1589.

22. Matarin M., Simon-Sanchez J., Fung H.-C., Scholz S., Gibbs J.R., Hernandez D.G., Crews C., Britton A., De Vrieze F.W., Brott T.G., et al. (2008). Structural genomic variation in ischemic stroke. Neurogenetics. 9, 101–108.

23. Keum S., Marchuk D.A. (2009). A locus mapping to mouse chromosome 7 determines infarct volume in a mouse model of ischemic stroke. Circ Cardiovasc Genet. 2, 591–598.

24. Keum S., Lee H.K., Chu P.-L., Kan M.J., Huang M.-N., Gallione C.J., Gunn M.D., Lo D.C., Marchuk D.A. (2013). Natural genetic variation of integrin alpha L (Itgal) modulates ischemic brain injury in stroke. PLoS Genet. 9, e1003807.

25. Chu P.-L., Keum S., Marchuk D.A. (2013). A novel genetic locus modulates infarct volume independently of the extent of collateral circulation. Physiol. Genomics. 45, 751–763.

26. Lee H.K., Keum S., Sheng H., Warner D.S., Lo D.C., Marchuk D.A. (2016). Natural allelic variation of the IL-21 receptor modulates ischemic stroke infarct volume. J Clin Invest. 126, 2827–2838.

27. Lee H.K., Koh S., Lo D.C., Marchuk D.A. (2018). Neuronal IL-4Rα modulates neuronal apoptosis and cell viability during the acute phases of cerebral ischemia. FEBS J. 285, 2785–2798.

28. Lee H.K., Widmayer S.J., Huang M.-N., Aylor D.L., Marchuk D.A. (2019). Novel neuroprotective loci modulating ischemic stroke volume in wild-derived inbred mouse strains. Genetics. 213, 1079–1092.

29. Lee H.K., Wetzel-Strong S.E., Aylor D.L., Marchuk D.A. (2021). A neuroprotective locus modulates ischemic stroke infarction independent of collateral vessel anatomy. Fornt. Neurosci. 15, 705160.

30. Wang S., Zhang H., Wiltshire T., Sealock R., Faber J.E. (2012). Genetic Dissection of the Canq1 Locus Governing Variation in Extent of the Collateral Circulation. PLoS One. 7, e31910.

31. Zhang H., Prabhakar P., Sealock R., Faber J.E. (2010). Wide genetic variation in the native pial collateral circulation is a major determinant of variation in severity of stroke. J. Cereb Blood Flow Metab. 30, 923–934.

32. Lucitti J.L., Sealock R., Buckley B.K., Zhang H., Xiao L., Dudley A.C., Faber J.E. (2016). Variants of Rab GTPase-Effector Binding Protein-2 Cause Variation in the Collateral Circulation and Severity of Stroke. Stroke. 47, 3022–3031.

33. Kofler N., Corti F., Rivera-Molina F., Deng Y., Toomre D., Simons M. (2018). The Rab-effector protein RABEP2 regulates endosomal trafficking to mediate vascular endothelial growth factor receptor-2 (VEGFR2)-dependent signaling. J Biol Chem. 293, 4805–4817.

34. Zhu G., Zhai P., Liu J., Terzyan S., Li G., Zhang X.C. (2004). Structural basis of Rab5-Rabaptin5 interaction in endocytosis. Nat Struct Mol Biol. 11, 975–983.

35. Jumper J., Evans R., Pritzel A., Green T., Figurnov M., Ronneberger O., Tunyasuvunakool K., Bates R., Žídek A., Potapenko A., et al. (2021). Highly accurate protein structure prediction with AlphaFold. Nature. 596, 583–589.

36. Varadi M., Anyango S., Deshpande M., Nair S., Natassia C., Yordanova G., Yuan D., Stroe O., Wood G., Laydon A., et al. (2021). AlphaFold Protein Structure Database: massively expanding the structural coverage of protein-sequence space with high-accuracy models. Nucleic Acids Res. 50, D439–D444.

37. Yan Y., Tao H., He J., Huang S.-Y. (2020). The HDOCK server for integrated protein–protein docking. Nat Protoc. 15, 1829–1852.

38. Gournier H., Stenmark H., Rybin V., Lippé R., Zerial M. (1998). Two distinct effectors of the small GTPase Rab5 cooperate in endocytic membrane fusion. EMBO J. 17, 1930–1940.

39. Hodge R.D., Bakken T.E., Miller J.A., Smith K.A., Barkan E.R., Graybuck L.T., Close J.L., Long B., Johansen N., Penn O., et al. (2019). Conserved cell types with divergent features in human versus mouse cortex. Nature. 573, 61–68.

40. O’Collins V.E., Macleod M.R., Donnan G.A., Horky L.L., van der Worp B.H., Howells D.W. (2006). 1,026 experimental treatments in acute stroke. Ann Neurol. 59, 467–477.

41. Ginsberg M.D. (2008). Neuroprotection for ischemic stroke: past, present and future. Neuropharmacology. 55, 363–389.

42. Plenge R.M., Scolnick E.M., Altshuler D. (2013). Validating therapeutic targets through human genetics. Nat Rev Drug Discov. 12, 581–594.

43. Nelson M.R., Tipney H., Painter J.L., Shen J., Nicoletti P., Shen Y., Floratos A., Sham P.C., Li M.J., Wang J., Cardon L.R., et al. (2015). The support of human genetic evidence for approved drug indications. Nat Genet. 47, 856–860.

44. King E.A., Davis J.W., Degner J.F. (2019). Are drug targets with genetic support twice as likely to be approved? Revised estimates of the impact of genetic support for drug mechanisms on the probability of drug approval. PLOS Genet. 15, e1008489.

45. Turcot V., Lu Y., Highland H.M., Schurmann C., Justice A.E., Fine R.S., Bradfield J.P., Esko T., Giri A., Graff M., et al. (2018). Protein-altering variants associated with body mass index implicate pathways that control energy intake and expenditure in obesity. Nat Genet. 50, 26–41.

46. Christakoudi S., Evangelou E., Riboli E., Tsilidis K.K. (2021). GWAS of allometric body-shape indices in UK Biobank identifies loci suggesting associations with morphogenesis, organogenesis, adrenal cell renewal and cancer. Sci Rep. 11, 10688.

47. Vujkovic M., Keaton J.M., Lynch J.A., Miller D.R., Zhou J., Tcheandjieu C., Huffman J.E., Assimes T.L., Lorenz K., Zhu X., et al. (2020). Discovery of 318 new risk loci for type 2 diabetes and related vascular outcomes among 1.4 million participants in a multi-ancestry meta-analysis. Nat Genet. 52, 680–691.

48. Berndt S.I., Gustafsson S., Mägi R., Ganna A., Wheeler E., Feitosa M.F., Justice A.E., Monda K.L., Croteau-Chonka D.C., Day F.R., et al. (2013). Genome-wide meta-analysis identifies 11 new loci for anthropometric traits and provides insights into genetic architecture. Nat Genet. 45, 501–12.

49. Grasby K.L., Jahanshad N., Painter J.N., Colodro-Conde L., Bralten J., Hibar D.P., Lind P.A., Pizzagalli F., Ching C.R.K., McMahon M.A.B., et al. (2020). The genetic architecture of the human cerebral cortex. Science. 367, eaay6690.

50. Nalls M.A., Blauwendraat C., Vallerga C.L., Heilbron K., Bandres-Ciga S., Chang D., Tan M., Kia D.A., Noyce A.J., Xue A., et al. (2019). Identification of novel risk loci, causal insights, and heritable risk for Parkinson’s disease: a meta-analysis of genome-wide association studies. Lancet Neurol. 18, 1091–1102.

51. Hill W.D., Marioni R.E., Maghzian O., Ritchie S.J., Hagenaars S.P., McIntosh A.M., Gale C.R., Davies G., Deary I.J. (2019). A combined analysis of genetically correlated traits identifies 187 loci and a role for neurogenesis and myelination in intelligence. Mol Psychiatry. 24, 169–181.

